# Timing of arrival in the breeding area is repeatable and affects reproductive success in a non-migratory population of blue tits

**DOI:** 10.1101/626788

**Authors:** Carol Gilsenan, Mihai Valcu, Bart Kempenaers

## Abstract

1. Events in one part of the annual cycle often affect the performance (and subsequently fitness) of individuals later in the season (carry-over effects). An important aspect of this relates to the timing of activities. For example, many studies on migratory birds have shown that relatively late spring arrival in the breeding area reduces both the likelihood of getting a mate or territory and reproductive success.
2. In contrast, relatively little is known about movements of individuals in non-migratory populations during the non-breeding season. Few studies have investigated the timing of arrival at the breeding area in such species, possibly due to the assumption that most individuals remain in the area during the non-breeding season.
3. In this study, we used four years of data from a transponder-based automated recording system set up in a non-migratory population of blue tits (*Cyanistes caeruleus*) to describe individual variation in arrival at the breeding site. We investigated whether this variation can be explained by individual characteristics (sex, body size, or status), and we assessed its effect on aspects of reproductive success in the subsequent breeding season.
4. We found substantial variation in arrival date and demonstrate that this trait is individual-specific (repeatable). Females arrived later than males, but the arrival dates of social pair members were more similar than expected by chance. Arrival predicted both whether an individual would end up breeding that season, and several aspects of its breeding success.
5. Our study suggests that non-migratory species show a form of incipient migration behaviour in that they leave the breeding area during the non-breeding season. We conclude that the timing of pre-breeding events, in particular arrival date, may be an overlooked, but important, fitness-relevant trait in non-migratory species.

## INTRODUCTION

Events and processes in one part of the annual cycle often affect the performance of individuals later in the season, a phenomenon known as “carry-over effects” (reviewed in Norris & Marra, 2007; Harrison, Blount, Inger, Norris, & Bearhop 2011). An important aspect of such carry-over effects relates to timing of behaviour. Indeed, the timing of activities during the non-breeding season, such as the timing of departure from the wintering area, may be linked to fitness in the subsequent breeding season.

In migratory birds, many studies have shown that males typically arrive in the breeding area earlier than females (reviewed in Morbey & Ydenberg, 2001), and that arriving relatively early is linked to fitness benefits for both males and females. Early-arriving individuals have a greater choice of territories and hence an increased likelihood of acquiring one of high quality, or indeed any territory at all (both sexes: Bensch & Hasselquist, 1991; males: Aebischer, Perrin, Krieg, Studer, & Meyer 1996), increased chances of obtaining a mate (Lozano, Perreault, & Lemon 1996; Currie, Thompson, & Burke 2000), and higher breeding success (Perrins, 1970; Forstmeier, 2002; Hötker, 2002; Norris, Marra, Kyser, Sherry, & Ratcliffe 2004; Vergara, Aguirre, & Fernández-Cruz 2007). In males, earlier arrival may also be related to higher extra-pair mating success (Alatalo, Carlson, Lundberg, & Ulfstrand 1981; Langefors, Hasselquist, & von Schantz 1998). Furthermore, arriving earlier allows a longer breeding season and may thus translate into more breeding attempts per season, either in species which produce multiple broods (Dunn & Møller, 2014), or in the context of renesting after failure of the first breeding attempt (Cooper, Murphy, Redmond, & Dolan 2011). Earlier arrival in the year of first return to a breeding site has also been linked to the probability of breeding in subsequent years in a long-lived seabird (Becker et al., 2008). These fitness benefits, however, do not necessarily lead to selection on early arrival, because they may be offset by survival costs. Individuals arriving early may face harsher environmental conditions, leading to higher risk of mortality (McNamara, Welham, & Houston 1998; Smith & Moore, 2003; Smith & Moore, 2005). This may lead to a relationship between individual quality and arrival date, if only high-quality individuals can withstand these suboptimal conditions (Kokko, 1999; Hatch, Smith, & Owen 2010).

For many migratory species, previous studies have reported that the timing of arrival at the breeding site is individually repeatable (van Wijk, Bauer, & Schaub 2016; Kentie et al., 2017; reviewed in Both, Bijlsma, & Ouwehand 2016). Several reasons have been proposed for this repeatability, including consistent individual differences in departure from the wintering grounds (Both et al., 2016), which can arise from individuals responding consistently to environmental cues (Petersen, 1992), and differences in individual morphology which affect migration speed (e.g. wing length; Both et al., 2016). Consistency in arrival (and/or departure from wintering grounds) may also reflect intrinsic differences in timing among individuals (Alerstam, Hake, & Kjellén 2006; Vardanis, Klaassen, Strandberg, & Alerstam 2011; Stanley, MacPherson, Fraser, McKinnon, & Stutchbury 2012; López-López, García-Ripollés, & Urios 2014), which in turn may arise from variation in individual quality, e.g. speed or stamina during sustained migratory flight (Both & Visser, 2001).

In non-migratory or year-round resident species, the effects of the timing of breeding (start of laying) on fitness have been well documented in many populations (e.g. Perrins & McCleery, 1989; Thorley & Lord, 2015; reviewed in Verhulst & Nilsson, 2008). However, the timing of arrival in the breeding area has not been considered previously, perhaps because it is assumed that most individuals do not leave the local area. Furthermore, data on the whereabouts of individuals outside the breeding season are often lacking. For example, in studies of post-breeding dispersal, i.e. movements from one breeding site to the next (e.g. Harvey, Greenwood, & Perrins 1979; Andreu & Barba, 2006; Valcu & Kempenaers, 2008), the timing of leaving from the first site and of arriving at the second site is often unknown.

Research on timing of arrival in non-migratory species has mostly focused on yearling individuals in the context of natal dispersal, i.e. movements from the site of fledging to the first breeding site (Greenwood, Harvey, & Perrins 1979; Lens & Dhondt, 1994; Verhulst, Perrins, & Riddington 1997; Ortego, García-Navas, Ferrer, & Sanz 2011; Pakanen, Koivula, Orell, Rytkönen, & Lahti 2016; reviewed in Greenwood & Harvey, 1982; Clarke, Sæther, & Røskaft 1997). For example, a recent study on yearling great tits (*Parus major*) reported that individuals grouped into winter flocks assortatively depending on arrival date in the population, whereby those that arrived later were less likely to breed that season (Farine & Sheldon, 2015). In the same population, immigrant females that arrived later in winter were also more likely to have a failed breeding attempt compared to female local recruits or early-arriving immigrants (Kidd, Sheldon, Simmonds, & Cole 2015).

We studied a non-migratory population of blue tits (*Cyanistes caeruleus*), and used an automated monitoring system at feeders and nestboxes to estimate the arrival date of all adult individuals in the breeding area. The main aims of our study were (a) to describe individual variation in arrival date, (b) to investigate whether arrival date is an individual-specific trait, (c) to explore if and how arrival date relates to pairing, timing of breeding, and measures of reproductive success, including extra-pair paternity, and (d) to investigate whether between-individual differences in timing of arrival can be explained by individual characteristics such as sex, body size, or status.

## MATERIALS AND METHODS

### Study species and site

The blue tit is a small passerine, present in large numbers across much of Europe. It is a secondary cavity-nester which readily accepts nestboxes for roosting and breeding. The majority of blue tits are socially monogamous, but extra-pair paternity is common (Schlicht, Valcu, & Kempenaers 2015). In our population, pairs produce one clutch per season, except when renesting after clutch or brood failure. The blue tit is a partial migrant in the northern part of its range (Nilsson, Lindström, Jonzén, Nilsson, & Karlsson 2006; Smallegange, Fiedler, Köppen, Geiter, & Barlein 2010), but is considered to be non-migratory in the rest of Europe (Stenning, 2018).

The study site in southern Germany (“Westerholz”, 48°08’26” N, 10°53’29” E) is a 40 ha protected part of a larger mixed deciduous/coniferous forest. Since 2007, the site has contained 277 nestboxes, and 60–170 pairs breed in these boxes each year. The data for this study were collected between August 2014 and July 2017. In October 2014, we installed 16 feeders around the edge of the study site, and filled them with chopped peanuts each year between October and March.

### General field procedures

To record breeding activities, we visited all nestboxes at least once a week between early March and early June. We recorded (1) the date of appearance of the first nesting material, (2) the date of nest completion (nest cup lined with feathers, hair, or other soft material), (3) the date of the first egg (hereafter “lay date”), based on daily checks of completed nests, (4) clutch size, and (5) the date the first egg(s) hatched (hereafter “hatch date”), based on daily nest checks around the estimated hatch date. When the nestlings were 14 days old, we ringed them, measured their tarsus (to the nearest 0.1 mm) and body mass (± 0.1 g), and took a blood sample (∼ 50 µl) by brachial venipuncture. When the nestlings were 19 days old, we checked the nest daily to determine fledging success and the date(s) of fledging.

Each year, from October/November to March, we checked nestboxes at least once per month at night, and trapped all roosting blue tits that did not carry a transponder (see below). We also caught individuals (1) with mist-nets at the feeder sites between October and March, (2) with snap-traps or mist-nets in their territory in March and April before nest completion (only individuals without a transponder), and (3) in their nestbox while they fed 9–11 day-old nestlings in May or June (only if not caught before).

Upon first capture, we ringed each individual with a metal ring and 1–3 plastic colour rings, measured tarsus (to the nearest 0.1 mm), primary wing feather length (± 0.5 mm) and body mass (± 0.1 g), scored age (yearling or adult; following Jenni & Winkler, 1994), and took a blood sample (∼ 20 µl) from the brachial vein. We also equipped each bird with a passive integrated transponder (PIT, BIOMARK HPT8 animal tag 134.2 kHz FDXB, 8.4 mm x 1.4 mm, 0.03 g, Biomark, Boise, ID, U.S.A.), which we inserted subcutaneously on the back (Schlicht & Kempenaers, 2015). All birds were released close to the location where they had been caught. For this study, we caught a total of 1,306 individuals.

### Nest and feeder monitoring system

All visits of PIT-tagged individuals to any nestbox or feeder were recorded continuously with an automated RFID-based monitoring system (see Schlicht, Girg, Loës, Valcu, & Kempenaers 2012 for details). Briefly, each nestbox and feeder was equipped with a transponder-reading device, a real-time clock, and a data storage device. Each nestbox also had two light barriers, one each at the outside and inside of the nest hole. Whenever a PIT-tagged bird landed at the nest hole or at the entrance to a feeder, its identity and the associated date and time were recorded. All visit data were stored on SD cards which we exchanged manually every 1–4 weeks. We checked all nestboxes and feeders regularly to ensure they were functioning properly and to change batteries when needed. In total, across all study years, the nestboxes registered 25,700,512 distinct records of 1,467 individuals, while the feeders registered 1,398,664 distinct records of 709 individuals.

### Parentage analysis

We extracted DNA from all blood samples and from all egg or nestling tissue samples, genotyped all samples using 14 highly polymorphic microsatellite markers and one sex chromosome-linked marker, and determined parentage following the procedures outlined in Schlicht et al. (2015). Because the identity of the biological mother is usually known with certainty, the probability of erroneous assignment of males as sires of the nestlings is close to zero.

### Data analysis

We defined a blue tit “season” as the period between 1 August in Year X and 31 July in Year X+1, based on the end of the breeding season in our population (mid-June) and the absence of nestbox visits in July.

For each individual, we defined its “arrival date” in the breeding area as the earliest record in a given season. To exclude biases arising from different methods of data collection or due to variation in the timing of capture, we only considered individuals that had been PIT-tagged in a previous season. We further included only data from automated recordings at feeders and nestboxes, excluding 22 individuals whose first record in a given season was a capture. We obtained data on arrival date for a total of 441 individuals; for 139 individuals, we recorded arrival date in at least two seasons. All sample sizes mentioned in the Results are for the number of distinct individuals, unless otherwise stated.

We analysed repeatability (*r*) in arrival date for both sexes combined and separately for individuals of different status: males and females, local recruits (individuals born in the study site), and breeders (individuals that were known to breed in a nestbox in the current season). Repeatability estimates are based on linear mixed-effect models (REML) for Gaussian traits, including only individuals with data for at least two seasons, and with individual identity as a random effect. For each repeatability estimate, we also estimated 95% confidence intervals (CI) by bootstrapping with 1,000 iterations.

We then assessed whether between-individual variation in arrival date can be explained by the individual characteristics sex, body size or status (local recruit or immigrant). We did this for males and females separately, and for two subgroups of individuals: those that bred in the study area in the current season, and those that were local recruits. For the latter, we tested whether fledging date predicted future arrival date. For these analyses, we used mixed-effect models with arrival date as the response variable, with a combination of the predictor variables mentioned above and with individual identity as a random effect.

We tested whether arrival date predicted whether an individual ended up breeding in the study area using a binomial generalised mixed-effect model with breeding status (breeder/non-breeder) as the response variable, and with arrival date as the predictor. For those individuals that did breed, we used two methods to test whether the arrival date of pair members were correlated. (1) For each of the three seasons, we calculated the Pearson’s correlation coefficient between the first-and the second-arriving pair member. (2) We used a linear mixed-effect model with arrival date of the second-arriving individual as the response variable and the arrival date of the first-arriving individual as the predictor. We included the sex of the later-arriving individual as a covariate in the mixed model, because its arrival date can be a result of its sex *per se*, and not necessarily linked to the arrival of its mate.

We also assessed whether arrival date explained variation in reproductive success. For males, we tested whether arrival date predicted paternity loss in their own brood and extra-pair siring success. We used binomial generalized linear mixed-effect models with the measure of reproductive success, i.e. paternity loss/gain (yes/no), as the response variable and with arrival date as the predictor. For females, we tested whether arrival date affected lay date, clutch size, and fledging success (number of chicks fledged). We used linear mixed-effect models with the measure of reproductive success as the response variable and with arrival date as the predictor. All models included individual identity as a random effect.

We used the free software R (version 3.4.1 and later; R Core Team, 2017) for statistical analyses. For repeatability estimates we used the “rptR” package (Stoffel, Nakagawa, & Schielzeth 2017) and for mixed model analyses the “lme4” package (Bates, Mächler, Bolker, & Walker 2015) with the package “multcomp” (Hothorn, Bretz, & Westfall 2008) to obtain *p*-values. Variables were scaled by subtracting the mean value for each year. Mixed-model estimates were plotted using the package “sjPlot” (Lüdecke, 2018).

## RESULTS

Individuals arrived at the breeding site between 1 August and 24 April (**Table 1; Fig. 1**). An individual’s mean arrival date correlated positively with the date of first capture in the study site in a previous season (Pearson’s correlation: *r* = 0.32, df = 439, *p* < 0.001), and with the date of second detection in the same season (linear mixed-effect model with second detection date as the response variable, arrival date as the predictor, and individual identity as a random effect: β_0_ ± SE (Intercept) = 2.47 ± 1.26; β ± SE = 38.65 ± 1.26, *z* = 30.71, *p* < 0.001, *n* = 405).

**Table 1.** Variation in arrival date of male and female blue tits in our study site for three seasons. Shown are mean and SE (in days), the first and last arrival date in that season, and the sample size (*n*).

| Category | Season | *n* | Mean arrival<br>date | SE | First arrival | Last arrival |
| --- | --- | --- | --- | --- | --- | --- |
| Males | 2014-2015 | 62 | 11 October | 10 | 2 August | 20 February |
|  | 2015-2016 | 104 | 24 October | 7 | 2 August | 25 March |
|  | 2016-2017 | 162 | 13 November | 8 | 1 August | 15 April |
| Females | 2014-2015 | 56 | 15 December | 14 | 2 August | 17 March |
|  | 2015-2016 | 97 | 13 December | 9 | 14 August | 27 March |
|  | 2016-2017 | 126 | 17 December | 9 | 1 August | 24 April |

**Figure 1.**
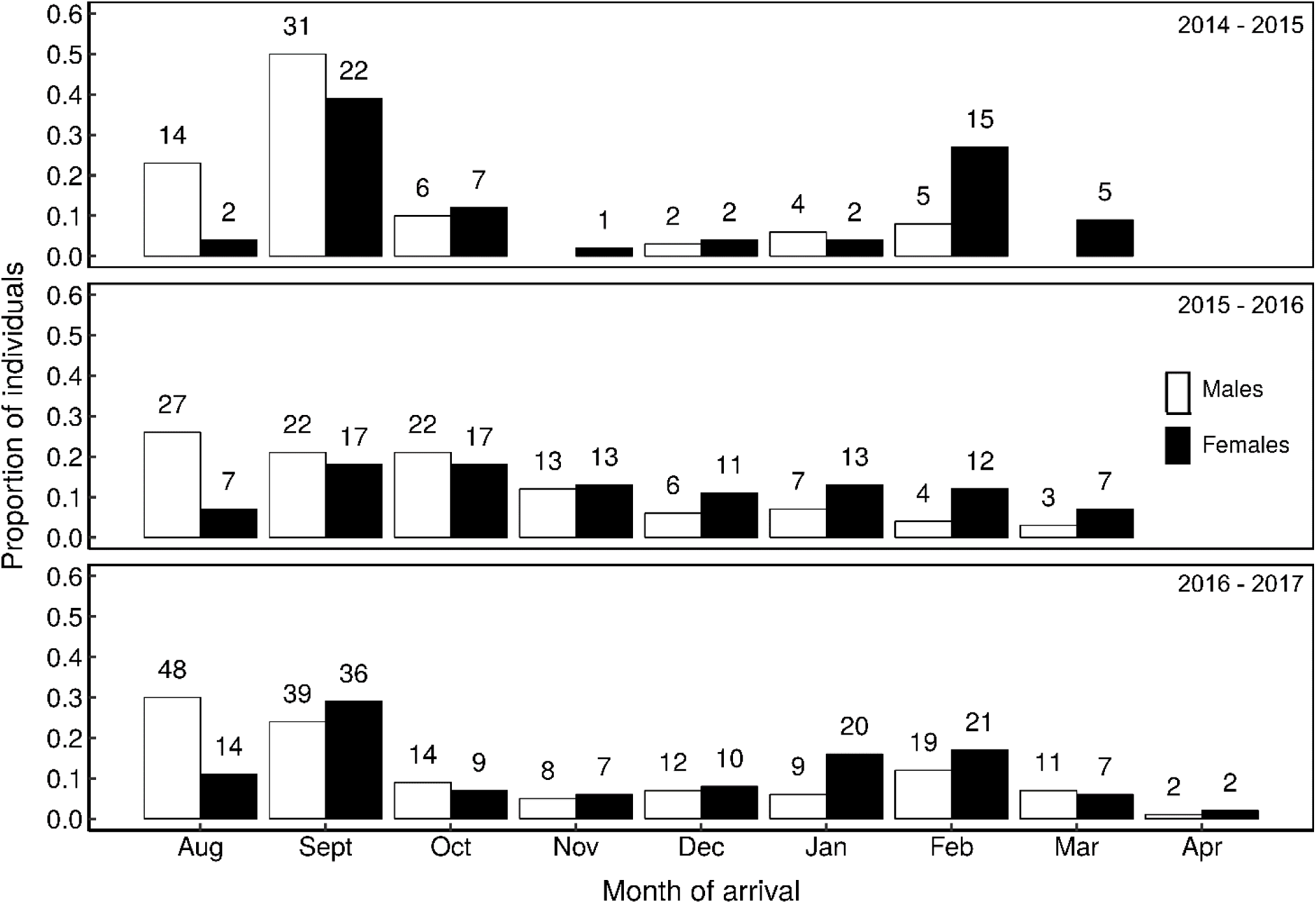
Frequency distribution of arrival dates of adult male (white bars) and female (black bars) blue tits in Westerholz, southern Germany, in three seasons (2014–2017). Shown are the proportions of the total number of males and females in a given season arriving each month. Numbers above each bar indicate the number of individuals in each group. Only individuals that had been caught and PIT-tagged in a previous season are included.

Arrival date was highly repeatable across all individuals (**Table 2; Fig. 2**). Repeatability estimates were similar for males and females, and were higher when only recruits or breeders were considered (**Table 2**). The difference in arrival date between seasons (the “return window”) varied between individuals, with a median of 25 days for males and 32.5 days for females (**Fig. 2**).

**Table 2.** Repeatability (*r*) in arrival date (first detection at a nestbox or feeder in the study area in a given season) for different categories of blue tits.

| Category | Model covariates* | $r$ | 95% CI | $p$ |
| --- | --- | --- | --- | --- |
| **All individuals** |  |  |  |  |
| Sexes combined | Sex | 0.60 | 0.50-0.70 | 0.001 |
| Males | Breeder, tarsus, recruit | 0.59 | 0.44-0.73 | 0.001 |
| Females | Breeder, tarsus, recruit | 0.57 | 0.41-0.73 | 0.001 |
| **All recruits** |  |  |  |  |
| Sexes combined | Sex, fledging date | 0.75 | 0.63-0.84 | 0.001 |
| Males | - | 0.79 | 0.65-0.88 | 0.001 |
| Females | - | 0.68 | 0.35-0.84 | 0.001 |
| **All breeders** |  |  |  |  |
| Sexes combined | Sex | 0.77 | 0.67-0.85 | 0.002 |
| Males | - | 0.88 | 0.78-0.93 | 0.001 |
| Females | - | 0.71 | 0.50-0.84 | 0.001 |

Sample sizes are as follows: males $n_{\text{total}} = 75$ , $n_{\text{recruit}} = 38$ , $n_{\text{immigrants}} = 37$ , $n_{\text{breeder}} = 56$ , $n_{\text{non-breeder}} = 40$ ; females $n_{\text{total}} = 64$ , $n_{\text{recruit}} = 20$ , $n_{\text{immigrants}} = 44$ , $n_{\text{breeder}} = 49$ , $n_{\text{non-breeder}} = 35$ .
\*Sex (male/female), breeder (yes/no), recruit (yes/no), fledging date (scaled per year).

**Figure 2.**
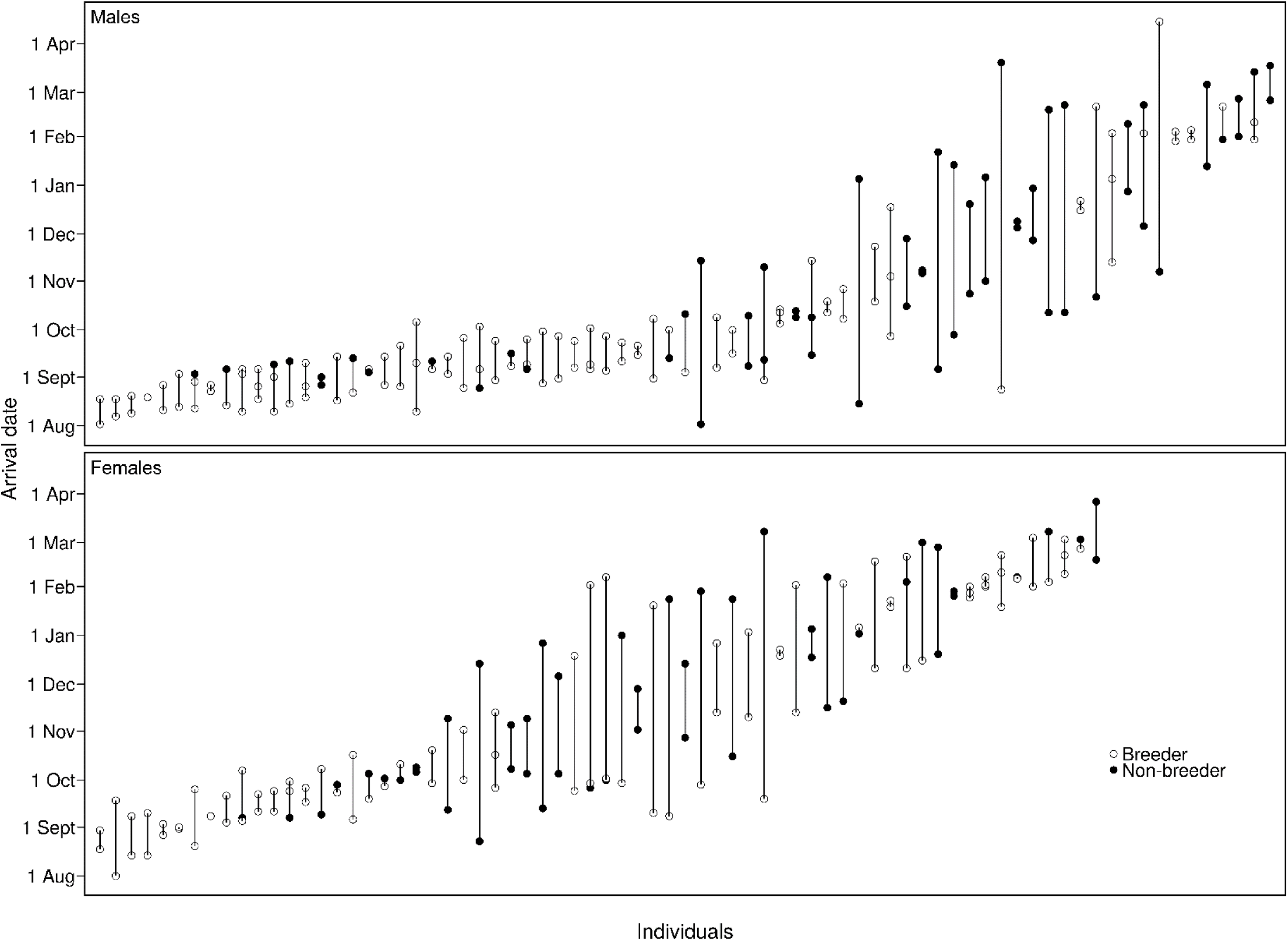
Arrival dates of adult male and female blue tits in the study site. Shown are all individuals recorded in at least two seasons (data from 2014–2017). Arrival dates of the same individual are connected by a line, and individuals are ordered by increasing mean arrival date from left to right. Open and filled circles indicate breeding and non-breeding in the study area in a given season respectively.

Females arrived later than males, both overall, and when only recruits or breeders were considered (**Table 3**). Arrival date did not depend on body size (tarsus length) or status (immigrant versus local recruit), neither in males, nor in females (**Table 3**), but individuals that ended up breeding in a given season arrived significantly earlier than non-breeders.

**Table 3.** Results of linear mixed models explaining variation in arrival date (first detection at a nestbox or feeder in the study site in a given season) for blue tits of different status.

| Category | Predictor* | Estimate ± SE | z value | p |
| --- | --- | --- | --- | --- |
| All individuals | *Intercept* | -0.29 ± 0.06 | | |
| | Sex | 0.44 ± 0.09 | 5.10 | <0.001 |
| Males | *Intercept* | 3.21 ± 2.03 | | |
| | Breeder | -0.34 ± 0.09 | -3.66 | <0.001 |
| | Tarsus | -0.20 ± 0.12 | -1.66 | 0.27 |
| | Recruit | 0.01 ± 0.11 | 0.10 | 1.00 |
| Females | *Intercept* | 1.57 ± 2.86 | | |
| | Breeder | -0.25 ± 0.11 | -2.25 | 0.072 |
| | Tarsus | -0.08 ± 0.17 | -0.48 | 0.95 |
| | Recruit | 0.19 ± 0.14 | 1.38 | 0.42 |
| Local recruits | *Intercept* | -0.27 ± 0.08 | | |
| | Sex | 0.53 ± 0.14 | 3.72 | <0.001 |
| | Fledging date | 0.02 ± 0.02 | 1.16 | 0.54 |
| Breeders | *Intercept* | -0.55 ± 0.08 | | |
| | Sex | 0.58 ± 0.12 | 5.03 | <0.001 |

Data from three seasons (2014-2017). Sample sizes are as follows: males $n_{\text{total}} = 237$ , $n_{\text{recruit}} = 130$ , $n_{\text{immigrants}} =$ 107, $n_{\text{breeder}} = 111$ , $n_{\text{non-breeder}} = 147$ ; females $n_{\text{total}} = 204$ , $n_{\text{recruit}} = 71$ , $n_{\text{immigrants}} = 133$ , $n_{\text{breeder}} = 108$ , $n_{\text{non-breeder}} =$ 116.
\* Sex (females relative to males), breeder (individuals that bred in that season relative to non-breeders), recruit (local recruits relative to immigrants), fledging date (scaled per year).

Arrival date predicted whether an individual would end up breeding that season, but the effect was only significant for males (**Fig. 3**).

**Figure 3.**
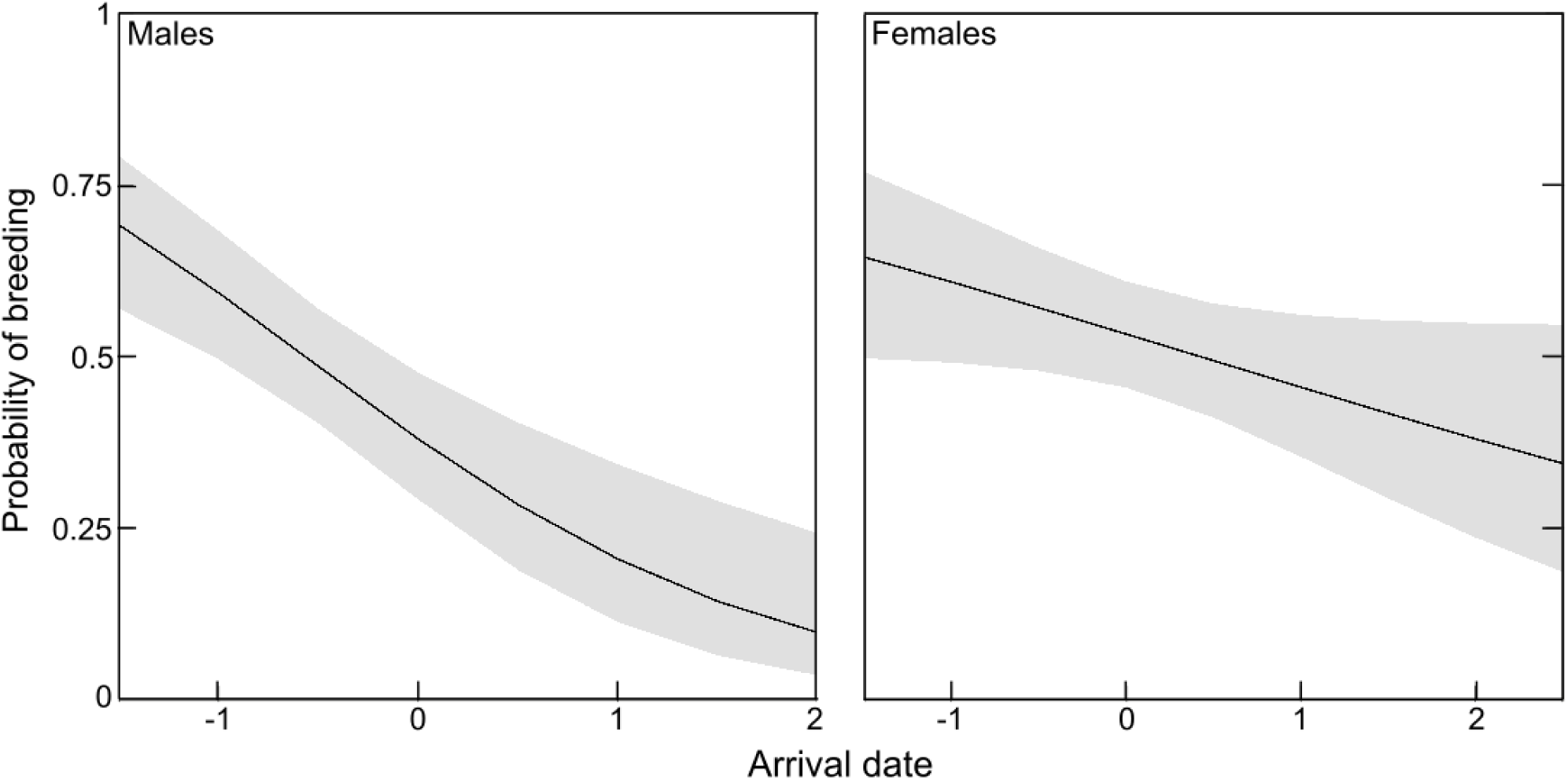
The predicted probability of breeding for male and female blue tits in relation to arrival date (scaled). Data are from three seasons (2014–2017). Shown are estimates with 95% confidence intervals from a generalized linear mixed model (logistic regression). Males: β_0_ ± SE = −0.49 ± 0.20, β ± SE = −0.87 ± 0.21, *z* = -4.13, *p* < 0.001, *n* = 237; females: β0 ± SE = 0.13 ± 0.16, β ± SE = −0.31 ± 0.16, *z* = −1.88, *p* = 0.12, *n* = 204.

The arrival date of pair members was positively correlated and was statistically significant in the two seasons with the highest sample size (Pearson’s correlation: *r*_*2014–2015*_ = 0.43, df = 13, *p* = 0.11; *r*_*2015–2016*_ = 0.46, df = 28, *p* = 0.01; *r*_*2016–2017*_ = 0.46, df = 42, *p* = 0.002). Individuals that bred together arrived in the study site relatively close in time to one another, even when the sex of the second (later) individual was taken into account (**Table 4; Fig. 4**).

**Table 4.**
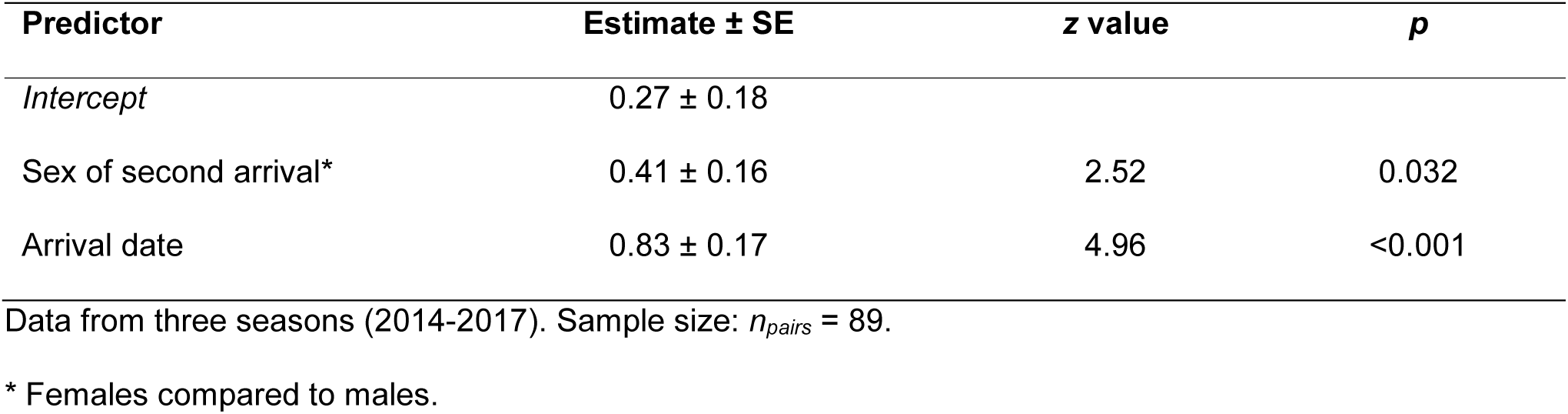
Results of a linear mixed model predicting the arrival date of the second (later) individual in a blue tit breeding pair by the arrival of the first (earlier) individual.

**Figure 4.**
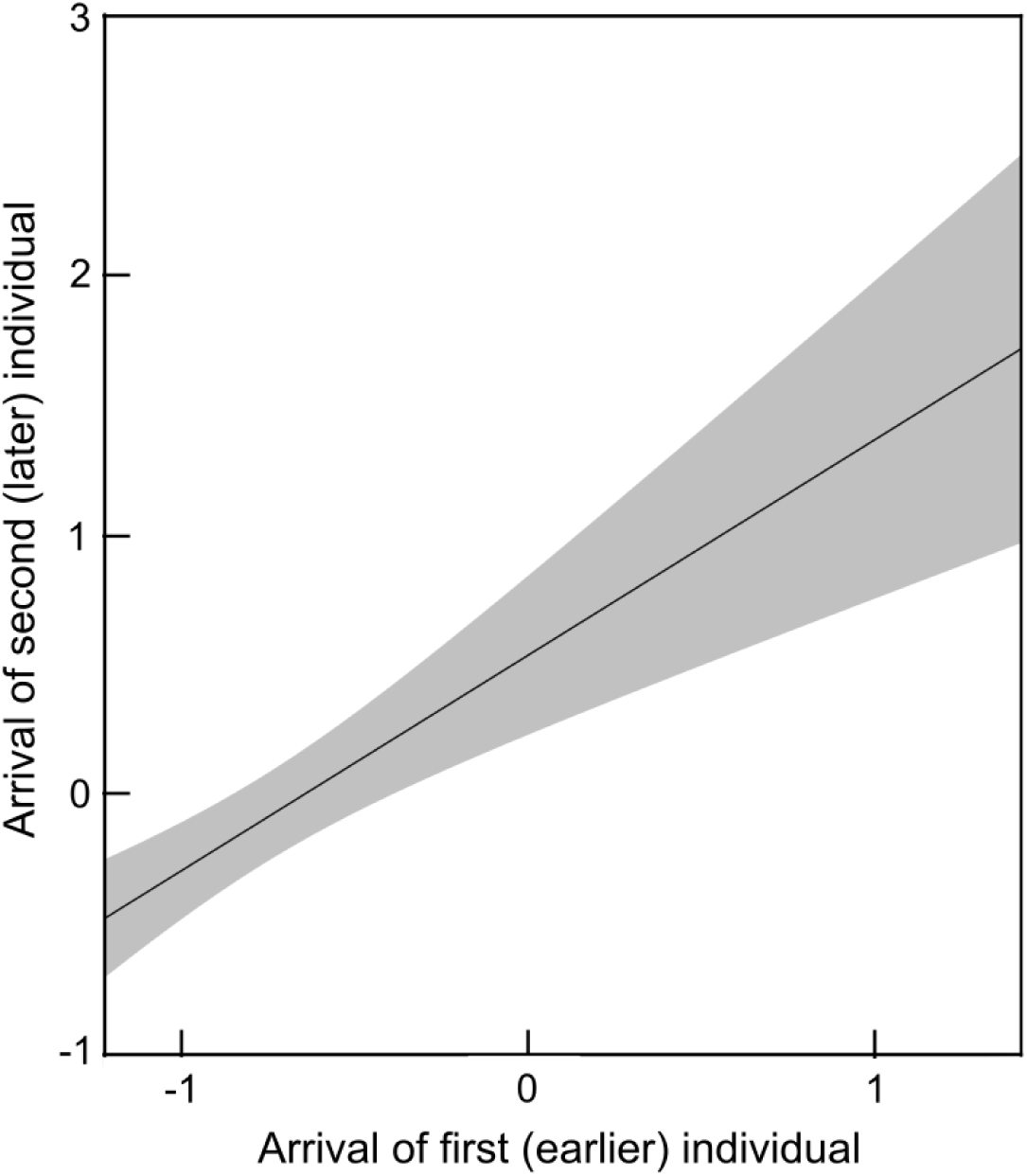
The relationship between arrival dates (scaled) of blue tit pair members in a particular season. Data are from three seasons (2014–2017). Shown are the model estimates with 95% confidence intervals.

Paternity loss, i.e. whether an extra-pair male sired at least one young in the focal male’s nest, was neither related to his arrival date, nor to that of his female mate (**Table 5**). However, males that arrived in the study area earlier in the season were more likely to sire extra-pair young during the breeding season, although this effect is not quite significant (**Table 5**).

**Table 5.** Results of linear mixed-effect models explaining variation in aspects of reproductive success of male and female blue tits.

| Category | Response* | Predictor* | Estimate ± SE | z value | p |
| --- | --- | --- | --- | --- | --- |
| Males | Paternity loss | *Intercept* | -0.18 ± 0.21 | | |
| | | Arrival date | -0.003 ± 0.23 | -0.01 | 1.00 |
| | Paternity gain | *Intercept* | -0.33 ± 0.23 | | |
| | | Arrival date | -0.55 ± 0.25 | -2.16 | 0.053 |
| Females | Mate's paternity loss | *Intercept* | -0.46 ± 0.20 | | |
| | | Arrival date | 0.05 ± 0.19 | 0.28 | 0.95 |
| | Lay date | *Intercept* | -0.04 ± 0.26 | | |
| | | Arrival date | 0.80 ± 0.26 | 3.05 | 0.005 |
| | Clutch size | *Intercept* | 0.02 ± 0.17 | | |
| | | Arrival date | -0.36 ± 0.17 | -2.20 | 0.055 |
| | Fledging success<sup>1</sup> | *Intercept* | 0.007 ± 0.027 | | |
| | | Arrival date | -0.13 ± 0.27 | -0.49 | 0.86 |

Data from three seasons (2014-2017). Sample sizes are as follows. Males paternity loss/gain: $n = 111$ ; paternity loss of females' social mates: $n_{females} = 108$ ; females breeding success (lay date, clutch size, fledging success): $n = 108$ .
\* All variables scaled per year, except paternity loss or gain (yes/no).
<sup>1</sup> Number of chicks fledged successfully (scaled).

For females that bred in the study area, arrival date predicted both lay date and clutch size, with later-arriving individuals laying later and laying smaller clutches, although the latter was not quite significant (**Table 5; Fig. 5**). There was no effect of female arrival date on subsequent fledging success (**Table 5**).

**Figure 5.**
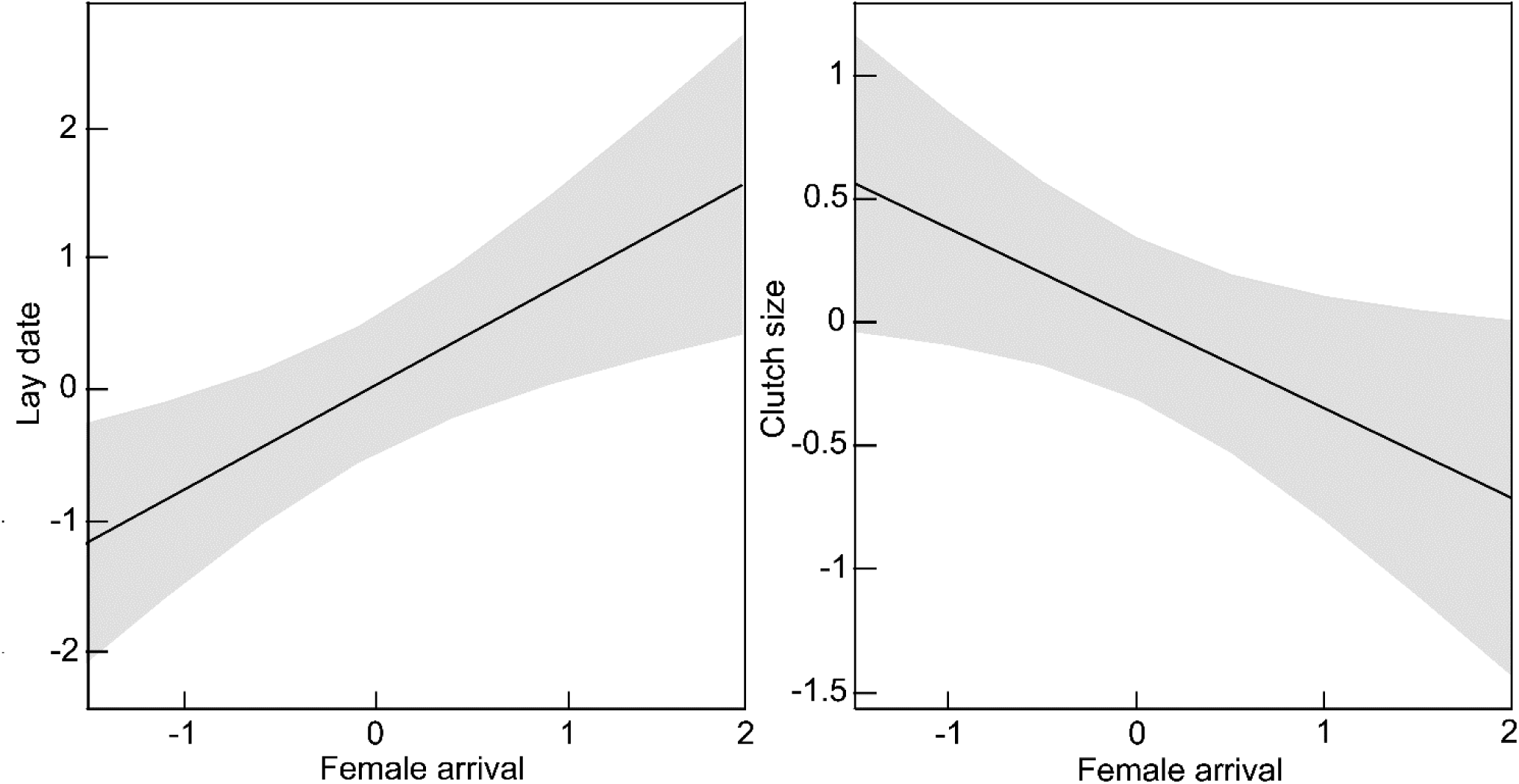
The relationship between female arrival date and her lay date (left panel) and clutch size (right panel). All variables are scaled within season. Data are from three seasons (2014–2017). Shown are model estimates with 95% confidence intervals (see Table 5 for model details).

## DISCUSSION

Our study shows substantial variation in arrival date in the breeding area for adult male and female blue tits from a non-migratory population. We further show that arrival date is a repeatable, individual-specific trait. Female blue tits arrived later than males, both overall, and when only breeding individuals or local recruits were considered, but arrival date of pair members was more similar than expected by chance. Arrival date predicted whether an individual would end up breeding that season (significant in males, non-significant trend in females), as well as aspects of male and female reproductive success. Taken together, our study suggests that the timing of pre-breeding events, in particular arrival date, is an important, fitness-relevant trait, even in a non-migratory species.

### Timing of arrival in migratory and non-migratory species

In migratory species, individuals show variation in their timing of arrival at the breeding site (Both et al., 2016). Our results suggest that this is also the case in a non-migratory species, whereby the variation in arrival date was even more pronounced. Our data stem from PIT-tagged individuals that were recorded automatically at nestboxes and feeders. Thus, individuals could still be present locally and not visiting these devices. However, previous studies suggest that it is more likely that they vanished from the local area for some time after the breeding season and returned after variable periods.

Although it is generally accepted that some adult tits remain resident in the breeding site over winter (Hinde, 1952; Gibb, 1954), Hinde (1953) commented in a study on great tits - another non-migratory, related species - that “some of the birds do show an increased tendency to wander during the migration seasons”. Another study on great tits showed that, based on comprehensive winter observations, breeding birds were more likely to remain resident in winter in urban parks than in forest habitats (Dhondt, Adriaensen, & Plompen 1996). Most forest-breeding birds are thought to leave the breeding area to spend the winter months in towns and villages (Gibb, 1954; Nilsson & Smith, 1988). Gibb (1954) noted that no tit species is exclusively sedentary, and that there are usually more blue and great tits breeding in the study site than numbers observed there in the middle of winter.

Blue and great tits, along with other passerine species, join mixed-species fission-fusion flocks during the non-breeding season (Ekman, 1989; see also review in Silk, Croft, Tregenza, & Bearhop 2014). Such flocks regularly travel straight-line distances of more than 3 km (Matechou, Cheng, Kidd, & Garroway 2015), which would already take them outside our study area (the longest distance between two locations, i.e. nestbox or feeder, within the study site is 954 m). Matechou et al. (2015) remarked that “the temporal patterns of these seasonal [flocking] movements are, at least superficially, similar to a partial migration between natural woodlands for breeding and external sites, likely towns and villages for overwintering” (see also references therein). The authors observed that individuals arrived back at the study site in a larger wave at the beginning of the non-breeding season, and in a smaller wave towards the end of the non-breeding season (see also Kokko, 1999), not unlike what we observed, although in our study arrival seems more continuous across the season (**Fig. 1**). We have no data on the movements of the Westerholz blue tits outside the study area during winter, but the aforementioned studies suggest that the assumption that most – if not all – individuals disperse from the study area for at least part of the non-breeding season is valid.

### Repeatability in timing of arrival

In migratory species, many studies have shown that timing of arrival at the breeding site is repeatable (Both et al., 2016). Proposed explanations include between-individual variation in wintering ground departure, intrinsic individual timing differences, and/or differences in individual quality. The results of our analyses indicate that the timing of arrival in the breeding area is also highly repeatable in blue tits. This suggests that “arrival” in the breeding area as estimated in this study for individuals that were caught in the previous seasons is indeed (close to) true arrival date.

In each season, the arrival dates of individuals spanned between eight and ten months (from August to between March and May, **Fig. 1**). Given the high repeatability of arrival date, this implies that there is substantial between-individual variation in this trait (see also Both et al., 2016). This suggests that some individuals either remained resident in the study site throughout winter, or chose to return from outside the breeding site early and long before breeding started. A winter translocation study on black-capped chickadees (*Poecile atricapillus*), another non-migratory species in the family Paridae, noted that “return [to the study area] was an individual response and not a flock behaviour pattern” (Odum, 1941*a*), suggesting that even in winter flocks, the propensity and timing of returning to the breeding area can differ markedly between individuals.

The date on which an individual was first caught was also positively correlated with its mean arrival date in the study area in later seasons, suggesting that capture date may be a useful proxy for arrival date. An individual’s arrival date (first detection) was also correlated with the date of second detection in the study area, further suggesting that first detection is a meaningful estimate of arrival in the study site.

Coupled with the finding that arrival times were correlated between pair members, the high repeatability indicates that individuals have distinctive pre-breeding schedules, and that arrival date may indeed be an important phenotypic trait, even in non-migratory species.

### Sex differences in timing of arrival

In migratory species, there is overwhelming evidence that males arrive at the breeding grounds before females (“protandry”, Rainio, Tøttrup, Lehikoinen, & Coppack 2007; reviewed in Morbey & Ydenberg, 2001; Kokko, Gunnarsson, Morrell, & Gill 2006). Several, non-mutually exclusive hypotheses have been proposed to explain protandry in such species, including (1) males choose to arrive early to avoid competition for territories or breeding sites with other males (given that males typically defend the territory), (2) selection favouring early-arriving males because they have a higher probability to breed or higher reproductive success, and (3) selection on females to arrive later to avoid suboptimal environmental conditions, or to reduce time awaiting males (reviewed in Morbey & Ydenberg, 2001).

In our non-migratory population of blue tits, we also found that males arrived earlier than females. Males, being dominant over females at feeding locations, may be more likely to remain resident in the breeding site over winter (Smith & Nilsson, 1987; Heldbjerg & Karlsson, 1997). Males may also be under selection to return to the breeding area – and to their former territory – as early as possible, because they may experience higher levels of competition for breeding resources, or even risk losing their territory, if they were to leave the area or return too late (“arrival-time hypothesis”; Ketterson & Nolan, 1976). Previous studies on parids suggest that individual patterns in winter residency may be linked to dominance (Sandell & Smith, 1991; Lahti, Koivula, Orell, & Rytkönen 1996; see also review in Ketterson & Nolan, 1983). Thus, subordinate individuals, such as females or lower-quality individuals, may be forced to leave the breeding area in winters with low levels of resources (food) because they are out-competed by dominant individuals (Smith & Nilsson, 1987; Heldbjerg & Karlsson, 1997). Indeed, a model predicts that if higher-quality individuals remain in the breeding site as year-round residents, as would be advantageous, then this may lead to the development of partial migration as a result of competition (Kokko, 1999).

### Timing of arrival and pair bonding

In populations of (partially) migratory species, it has been shown that individuals mate assortatively based on their arrival time in the breeding area (Village, 1985; Bearhop et al., 2005; Ludwig & Becker, 2008; see also Lozano et al., 1996) which may also be linked to age-dependent arrival (Coulson, 1966). Similarly, in this study we found that the arrival dates of future pair members were positively correlated (see also Lourenço et al., 2011). Colquhoun (1942) witnessed male blue tits performing pair-bonding displays to females from November into early spring, suggesting that blue tits may (re)establish pair bonds at variable times and often long before breeding starts (see also Odum, 1941*b*; Psorakis, Roberts, Rezek, & Sheldon 2012).

In our population, the difference in arrival date of former pair members also explained the occurrence of divorce. If the former pair members did not arrive within a short period, the probability of divorce strongly increased (Gilsenan, Valcu, & Kempenaers 2017). There are two potential processes that may explain this. One possibility is that former pair members typically stay together throughout winter (see also Psorakis et al., 2012). In that case, divorce may result from a decision or a process whereby the male and the female become separated, for example, because they join different winter flocks, or because one of them remains resident while the other individual leaves the breeding area. Alternatively, in blue tits, breeding pairs may not maintain their pair bond in between breeding seasons, i.e. they may not typically spend the winter together, but will breed together again if they both come back to the same area (the former territory) around the same time. In a short-lived species with high adult mortality such as the blue tit, individuals that would decide to wait for their former partner to return may risk not breeding at all, and hence may choose to pair up at the first opportunity with a different individual if their former mate is not (yet) around. Individual consistency in timing of arrival may thus be a strategy allowing individuals to breed with the same partner over multiple seasons (see also Gunnarsson, Gill, Sigurbjörnsson, & Sutherland 2004).

### Timing of arrival as a predictor of breeding success

The link between timing of arrival at the breeding area and the timing and success of breeding has been studied extensively in migratory species (e.g. Cristol, 1995; Aebischer,et al., 1996; Tryjanowski, Sparks, Ptaszyk, & Kosicki 2004; Gienapp & Bregnballe, 2012). Many of these studies have shown that the arrival schedules of individuals have far-reaching implications (“domino effect”, Piersma, 1987; e.g. Tarka, Hansson, & Hasselquist 2015; see also review in Alerstam, 2011). In migratory species, arrival date predicts when individuals breed (e.g. Bêty, Gauthier, & Giroux 2003), the size of the pool of potential males or territories they can choose from (“rank advantage hypothesis”, see Morbey & Ydenberg, 2001; Gunnarsson et al., 2006), and a multitude of breeding parameters (see Introduction). In general, individuals that arrive earlier are of better quality and have higher reproductive success (see review in Kokko, 1999).

In our study, individuals that arrived earlier had a higher probability of ending up breeding in the same season (significant in males, non-significant tendency in females; see also Le Bohec et al., 2007; Catry, Dias, Phillips, & Granadeiro 2013; Farine & Sheldon, 2015; Matechou et al., 2015). This suggests that a carry-over effect exists in our population, and that this effect may be sex-dependent (see also Saino et al., 2017). Due to competition for territories, nest sites or mates, males may be more constrained in their arrival timing. Indeed, we found that males of each status (overall, recruit, breeder) were somewhat more repeatable in their timing of arrival than females (**Table 2**). Variation in arrival date may also indicate differences in individual quality (since higher-quality individuals are expected to better endure costs of arriving early; Kokko, 1999; Matechou et al., 2015) and lead to assortative mating for quality, since individual timing predicts mating (Farine & Sheldon, 2015; this study). Although we found no differences in arrival related to body size (measured as tarsus length, **Table 3**), this may not be the best measure of “quality” (Wilson & Nussey, 2010; see also Nilsson & Smith, 1988). Early-arriving males may have had more time (1) to find an unoccupied territory (possibly their former territory) or a mate, (2) to settle within the social hierarchy of the local population, or (3) to gain information about conspecifics’ spatiotemporal movements. The latter two may also help increase mating prospects or win competitive interactions (Adriaensen & Dhondt, 1990; Bai, Severinghaus, & Philippart 2012; Matechou et al., 2015).

Earlier arrival also increased the probability that a male sired extra-pair offspring (see also Langefors et al., 1998; Cooper et al., 2011). In blue tits (and other species), yearling males are less likely to sire extra-pair young than adult males (e.g. Schlicht et al., 2015). However, this cannot explain the current result, because all individuals in this study are adults. Early-arriving males may either be of higher quality and females may actively have chosen them for extra-pair copulations (Kempenaers et al., 1992; Langefors et al., 1998), or they may have had more time to interact with opposite-sex conspecifics than late-arriving males, which may influence their social status in the subsequent breeding season (e.g. Firth & Sheldon, 2016). However, whether interactions during the non-breeding season explain patterns of extra-pair paternity remains to be shown (Maldonado-Chaparro, Montiglio, Forstmeier, Kempenaers, & Farine 2018).

For female blue tits, later arrival was associated with later laying (see also Smith & Moore, 2005) and with a smaller clutch (**Fig. 5**). Breeding success usually decreases with lay date, not only due to differences in individual timing *per se* (“timing hypothesis”, Williams, 2012), but also because of differences in individual quality related to timing (“quality hypothesis”, Williams, 2012). Early arrival as well as early laying may be in itself an indicator of individual quality (see also Blums, Nichols, Hines, Lindberg, & Mednis 2005), for example reflecting variation in individual condition or in capability to gather resources (Nussey, Postma, Gienapp, & Visser 2005). A later start of laying often has knock-on effects on other breeding parameters including clutch size and fledging success (Perrins, 1979; Verhulst & Tinbergen, 1991), although we did not find evidence for an effect on fledging success in our study.

Of all individuals included in this study, 70% of males and 72% of females had bred in the study area the previous season, and these individuals also arrived earlier than those that had not bred (see also Lanctot, Sandercock, & Kempenaers 2000). Individuals which bred previously in the breeding area have more experience relative to those that did not, and in that sense may be “higher quality” (see also Smith, 1995). In any case, because all individuals included in our study were adults (after-second year), we can exclude that differences in male and female reproductive success between early-and late-arriving individuals were due to age effects (as age effects are mostly observed between yearling and older birds).

### Conclusions and suggestions for future work

In conclusion, migratory and non-migratory species may be more similar than previously appreciated, in the sense that individuals leave the breeding area to spend part of the non-breeding season elsewhere before returning to the same or a nearby breeding location. The main difference then lies in the scale of those movements. We suggest that it may be heuristically worthwhile to consider movements of “resident species” as incipient migratory behaviour, and to consider the timing of arrival as an important life-history trait. Further study of the causes and consequences of within-and between-individual variation in the timing of arrival is warranted. For example, we here show that arrival date in blue tits is repeatable, and it may thus be interesting to assess whether the trait is also heritable. We also need information about the whereabouts and the movement patterns of individuals outside the period when they are detected in the breeding area.

## ACKNOWLEDGEMENTS

We thank all field team members for helping to catch, process, and record data on the blue tits, especially Agnes Türk and Andrea Wittenzellner. We are grateful to Peter Loës and Peter Skripsky for their continued maintenance of the “smart nestboxes” and “smart feeders”, and to Alexander Girg and Sylvia Kuhn for genotyping. CG is a member of the International Max Planck Research School for Organismal Biology. The study was funded by the Max Planck Society (to BK).

## AUTHORS’ CONTRIBUTIONS

CG and BK conceived the study and devised the methodology. CG and MV analysed the data with input from BK, and CG and BK wrote the manuscript with input from MV. All authors approved the manuscript for publication.

## DATA ACCESSIBILITY

All data will be uploaded and made available on Dryad.

## REFERENCES

Adriaensen, F., & Dhondt, A. A. (1990). Population dynamics and partial migration of the European robin (*Erithacus rubecula*) in different habitats. Journal of Animal Ecology, 59(3), 1077–1090. doi: 10.2307/5033

Aebischer, A., Perrin, N., Krieg, M., Studer, J., & Meyer, D. R. (1996). The role of territory choice, mate choice and arrival date on breeding success in the Savi’s warbler *Locustella luscinioides*. Journal of Avian Biology, 27, 143–152. doi: 10.2307/3677143

Alatalo, R. V., Carlson, A., Lundberg, A., & Ulfstrand, S. (1981). The conflict between male polygamy and female monogamy: the case of the Pied Flycatcher *Ficedula hypoleuca*. The American Naturalist, 117, 738–753. doi: 10.1086/283756

Alerstam, T. (2011). Optimal bird migration revisited. Journal of Ornithology, 152 (Supplement I), S5–S23. doi: 10.1007/s10336-011-0694-1

Alerstam, T., Hake, M., & Kjellén, N. (2006). Temporal and spatial patterns of repeated migratory journeys by ospreys. Animal Behaviour, 71(3), 555–566. doi: 10.1016/j.anbehav.2005.05.016

Andreu, J., & Barba, E. (2006). Breeding dispersal of great tits *Parus major* in a homogeneous habitat: effects of sex, age, and mating status. Ardea, 94(1), 45–58.

Bai, M. -L., Severinghaus, L. L., & Philippart, M. T. (2012). Mechanisms underlying small-scale partial migration of a subtropical owl. Behavioral Ecology, 23(1), 153–159. doi: 10.1093/beheco/arr168

Bates, D., Mächler, M., Bolker, B., & Walker, S. (2015). Fitting linear mixed-effects models using lme4. Journal of Statistical Software, 67(1), 1–48. doi: 10.18637/jss.v067.i01

Bearhop, S., Fiedler, W., Furness, R. W., Votier, S. C., Waldron, S., Newton, J., … Farnsworth, K. (2005). Assortative mating as a mechanism for rapid evolution of a migratory divide. Science, 310(5747), 502–504. doi: 10.1126/science.1115661

Becker, P. H., Dittmann, T., Ludwigs, J. D., Limmer, B., Ludwig, S. C., Bauch, C., … Wendeln, H. (2008). Timing of initial arrival at the breeding site predicts age at first reproduction in a long-lived migratory bird. Proceedings of the National Academy of Sciences of the United States of America, 105(34), 12349–12352. doi: 10.1073/pnas.0804179105

Bensch, S., & Hasselquist, D. (1991). Territory infidelity in the polygynous great reed warbler *Acrocephalus arundinaceus*: the effect of variation in territory attractiveness. Journal of Animal Ecology, 60(3), 857–871. doi: 10.2307/5418

Bêty, J., Gauthier, G., & Giroux, J. (2003). Body condition, migration, and timing of reproduction in snow geese: a test of the condition-dependent model of optimal clutch size. The American Naturalist, 162(1), 110–121. doi: 10.1086/375680

Blums, P., Nichols, J. D., Hines, J. E., Lindberg, M. S., & Mednis, A. (2005). Individual quality, survival variation and patterns of phenotypic selection on body condition and timing of nesting in birds. Oecologia, 143(3), 365–376. doi: 10.1007/s00442-004-1794-x

Both, C., Bijlsma, R. G., & Ouwehand, J. (2016). Repeatability in spring arrival dates in pied flycatchers varies among years and sexes. Ardea, 104(1), 3–21. doi: 10.5253/arde.v104i1.a1

Both, C., & Visser, M. E. (2001). Adjustment to climate change is constrained by arrival date in a long-distance migratory bird. Nature, 411, 296–298. doi: 10.1038/35077063

Catry, P., Dias, M. P., Phillips, R. A., & Granadeiro, J. P. (2013). Carry-over effects from breeding modulate the annual cycle of a long-distance migrant: an experimental demonstration. Ecology, 94(6), 1230–1235. doi: 10.1890/12-2177.1

Clarke, A. L., Sæther, B. -E., & Røskaft, E. (1997). Sex biases in avian dispersal: a reappraisal. Oikos, 79(3), 429–438. doi: 10.2307/3546885

Colquhoun, M. K. (1942). Notes on the social behaviour of blue tits. British Birds, 35(11), 234–240.

Cooper, N. W., Murphy, M. T., Redmond, L. J., & Dolan, A. C. (2011). Reproductive correlates of spring arrival date in the Eastern Kingbird *Tyrannus tyrannus*. Journal of Ornithology, 152(1), 143–152. doi: 10.1007/s10336-010-0559-z

Coulson, J. C. (1966). The influence of the pair-bond and age on the breeding biology of the kittiwake gull *Rissa tridactyla*. Journal of Animal Ecology, 35(2), 269–279. doi: 10.2307/2394

Cristol, D. A. (1995). Early arrival, initiation of nesting, and social status: an experimental study of breeding female red-winged blackbirds. Behavioral Ecology, 6(1): 87–93. doi: 10.1093/beheco/6.1.87

Currie, D., Thompson, D. B. A., & Burke, T. (2000). Patterns of territory settlement and consequences for breeding success in the Northern wheatear *Oenanthe oenanthe*. Ibis, 142(3), 389–398. doi: 10.1111/j.1474-919X.2000.tb04435.x

Dhondt, A. A., Adriaensen, F., & Plompen, W. (1996). Between- and within-population variation in mate fidelity in the great tit. In J. M. Black (Ed.), Partnerships in Birds: The Study of Monogamy (pp. 235–248). Oxford, U.K.: Oxford University Press.

Dunn, P. O., & Møller, A. P. (2014). Changes in breeding phenology and population size of birds. Journal of Animal Ecology, 83(3), 729–739. doi: 10.1111/1365-2656.12162

Ekman, J. (1989). Ecology of non-breeding social systems of *Parus*. The Wilson Bulletin, 101(2), 263–288.

Farine, D. R., & Sheldon, B. C. (2015). Selection for territory acquisition is modulated by social network structure in a wild songbird. Journal of Evolutionary Biology, 28(3), 547–556. doi: 10.1111/jeb.12587

Firth, J. A., & Sheldon, B. C. (2016). Social carry-over effects underpin trans-seasonally linked structure in a wild bird population. Ecology Letters, 19(11), 1324–1332. doi: 10.1111/ele.12669

Forstmeier, W. (2002). Benefits of early arrival at breeding grounds vary between males. Journal of Animal Ecology, 71(1), 1–9. doi: 10.1046/j.0021-8790.2001.00569.x

Gibb, J. (1954). Population changes of titmice, 1947–1951. Bird Study, 1(2), 40–48. doi: 10.1080/00063655409475788

Gienapp, P., & Bregnballe, T. (2012). Fitness consequences of timing of migration and breeding in cormorants. PloS One, 7(9), e46165. doi: 10.1371/journal.pone.0046165

Gilsenan. C., Valcu, M., & Kempenaers, B. (2017). Difference in arrival date at the breeding site between former pair members predicts divorce in blue tits. Animal Behaviour, 133, 57–72. doi: 10.1016/j.anbehav.2017.09.004

Greenwood, P. J., & Harvey, P. H. (1982). The natal and breeding dispersal of birds. Annual Review of Ecology, Evolution, and Systematics, 13, 1–21. doi: 10.1146/annurev.es.13.110182.000245

Greenwood, P. J., Harvey, P. H., & Perrins, C. M. (1979). The role of dispersal in the great tit (*Parus major*): the causes, consequences and heritability of natal dispersal. Journal of Animal Ecology, 48(1), 123–142. doi: 10.2307/4105

Gunnarsson, T. G., Gill, J. A., Atkinson, P. W., Gélinaud, G., Potts, P. M., Croger, R. E., … Sutherland, W. J. (2006). Population-scale drivers of individual arrival times in migratory birds. Journal of Animal Ecology, 75(5), 1119–1127. doi: 10.1111/j.1365-2656.2006.01131.x

Gunnarsson, T. G., Gill, J. A., Sigurbjörnsson, T., & Sutherland, W. J. (2004). Arrival synchrony in migratory birds. Nature, 431, 646. doi: 10.1038/431646a

Harrison, X. A., Blount, J. D., Inger, R., Norris, D. R., & Bearhop, S. (2011). Carry-over effects as drivers of fitness differences in animals. Journal of Animal Ecology, 80(1), 4–18. doi: 10.1111/j.1365-2656.2010.01740.x

Harvey, P. H., Greenwood, P. J., & Perrins, C. M. (1979). Breeding area fidelity of great tits (*Parus major*). Journal of Animal Ecology, 48(1), 305–313. doi: 10.2307/4115

Hatch, M. I., Smith, R. J., & Owen, J. C. (2010). Arrival timing and haematological parameters in gray catbirds (*Dumetella carolinensis*). Journal of Ornithology, 151(3), 545–552. doi: 10.1007/s10336-009-0487-y

Heldbjerg, H., & Karlsson, L. (1997). Autumn migration of blue tits *Parus caeruleus* at Falsterbo, Sweden 1980-94: population changes, migration patterns and recovery analysis. Ornis Svecica, 7, 149–167.

Hinde, R. A. (1952). The behaviour of the great tit (Parus major) and some other related species. E.J. Brill, Leiden.

Hinde, R. A. (1953). Appetitive behaviour, consummatory act, and the hierarchical organisation of behaviour – with special reference to the great tit (*Parus major l*.). Behaviour, 5(1), 189–224. doi: 10.1163/156853953X00113

Hothorn, T., Bretz, F., & Westfall, P. (2008). Simultaneous inference in general parametric models. Biometrical Journal, 50(3), 346–363. doi: 10.1002/bimj.200810425

Hötker, H. (2002). Arrival of pied avocets *Recurvirostra avosetta* at the breeding site: effects of winter quarters and consequences for reproductive success. Ardea, 90(3), 379–387.

Jenni, L., & Winkler, R. (1994). Moult and Ageing of European Passerines. Academic Press, London.

Kempenaers, B., Verheyen, G. R., Van den Broeck, M., Burke, T., Van Broeckhoven, C., & Dhondt, A. (1992). Extra-pair paternity results from female preference for high-quality males in the blue tit. Nature, 357, 494–496. doi: 10.1038/357494a0

Kentie, R., Marquez-Ferrando, R., Figuerola, J., Gangoso, L., Hooijmeijer, J. C. E. W., Loonstra, A. H. J., … Piersma, T. (2017). Does wintering north or south of the Sahara correlate with timing and breeding performance in black-tailed godwits? Ecology and Evolution, 7(8), 2812–2820. doi: 10.1002/ece3.2879

Ketterson, E. D., & Nolan, V. (1976). Geographic variation and its climatic correlates in the sex ratio of Eastern-wintering dark-eyed juncos (*Junco hyemalis hyemalis*). Ecology, 57(4), 679–693. doi: 10.2307/1936182

Ketterson, E. D., & Nolan, V. (1983). The evolution of differential bird migration. Current Ornithology (Ed. R.F. Johnston), Vol 1. Springer, Boston, MA. doi: 10.1007/978-1-4615-6781-3_12

Kidd, L. R., Sheldon, B. C., Simmonds, E. G., & Cole, E. F. (2015). Who escapes detection? Quantifying the causes and consequences of sampling biases in a long-term field study. Journal of Animal Ecology, 84(6), 1520–1529. doi: 10.1111/1365-2656.12411

Kokko, H. (1999). Competition for early arrival in migratory birds. Journal of Animal Ecology, 68(5), 940–950. doi: 10.1046/j.1365-2656.1999.00343.x

Kokko, H., Gunnarsson, T. G., Morrell, L. J., & Gill, J. A. (2006). Why do female migratory birds arrive later than males? Journal of Animal Ecology, 75(6), 1293–1303. doi: 10.1111/j.1365-2656.2006.01151.x

Lahti, K., Koivula, K., Orell, M., & Rytkönen, S. (1996). Social dominance in free-living willow tits *Parus montanus*: determinants and some implications of hierarchy. Ibis, 138(3), 539–544. doi: 10.1111/j.1474-919X.1996.tb08075.x

Lanctot, R. B., Sandercock, B. K., & Kempenaers, B. (2000). Do male breeding displays function to attract mates or defend territories? The explanatory role of mate and site fidelity. Waterbirds: The International Journal of Waterbird Biology, 23(2), 155–164.

Langefors, Å., Hasselquist, D., & von Schantz, T. (1998). Extra-pair fertilizations in the sedge warbler. Journal of Avian Biology, 29(2), 134–144. doi: 10.2307/3677191

Le Bohec, C., Gauthier-Clerc, M., Grémillet, D., Pradel, R., Béchet, A., Gendner, J., & Le Maho, Y. (2007). Population dynamics in a long-lived seabird: I. Impact of breeding activity on survival and breeding probability in unbanded king penguins. Journal of Animal Ecology, 76(6), 1149–1160. doi: 10.1111/j.1365-2656.2007.01268.x

Lens, L., & Dhondt, A. A. (1994). Effects of habitat fragmentation on the timing of crested tit *Parus cristatus* natal dispersal. Ibis, 136(2), 147–152. doi: 10.1111/j.1474-919X.1994.tb01078.x

López-López, P., García-Ripollés, C., & Urios, V. (2014). Individual repeatability in timing and spatial flexibility of migration routes of trans-Saharan migratory raptors. Current Zoology, 60(5), 642–652. doi: 10.1093/czoolo/60.5.642

Lourenço, P. M., Kentie, R., Schroeder, J., Groen, N. M., Hooijmeijer, J. C. E. W., & Piersma, T. (2011). Repeatable timing of northward departure, arrival and breeding in black-tailed godwits *Limosa l. limosa*, but no domino effects. Journal of Ornithology, 152(4), 1023–1032. doi: 10.1007/s10336-011-0692-3

Lozano, G. A., Perreault, S., & Lemon, R. E. (1996). Age, arrival date and reproductive success of male American redstarts *Setophaga ruticilla*. Journal of Avian Biology, 27(2), 164–170. doi: 10.2307/3677146

Ludwig, S. C., & Becker, P. H. (2008). Supply and demand: causes and consequences of assortative mating in common terns *Sterna hirundo*. Behavioral Ecology and Sociobiology, 62, 1601–1611. doi: 10.1007/s00265-008-0589-1

Lüdecke, D. (2018). sjPlot: Data visualization for statistics in social science. R package version 2.6.1, URL https://CRAN.R-project.org/package=sjPlot. doi: 10.5281/zenodo.1308157

Maldonado-Chaparro, A. A., Montiglio, P. –O., Forstmeier, W., Kempenaers, B., & Farine, D. R. (2018). Linking the fine-scale social environment to mating decisions: a future direction for the study of extra-pair paternity. Biological Reviews, 93(3), 1558–1577. doi: 10.1111/brv.12408

Matechou, E., Cheng, S. C., Kidd, L. R., & Garroway, C. J. (2015). Reproductive consequences of the timing of seasonal movements in a nonmigratory wild bird population. Ecology, 96(6), 1641–1649. doi: 10.1890/14-0886.1

Morbey, Y. E., & Ydenberg, R. C. (2001). Protandrous arrival timing to breeding areas: a review. Ecology Letters, 4(6), 663–673. doi: 10.1046/j.1461-0248.2001.00265.x

McNamara, J. M., Welham, R. K., & Houston, A. I. (1998). The timing of migration within the context of an annual routine. Journal of Avian Biology, 29(4), 416–423. doi: 10.2307/3677160

Nilsson, A. L. K., Lindström, Å., Jonzén, N., Nilsson, S. G., & Karlsson, L. (2006). The effect of climate change on partial migration – the blue tit paradox. Global Change Biology, 12(10), 2014–2022. doi: 10.1111/j.1365-2486.2006.01237.x

Nilsson, J. A., & Smith, H. G. (1988). Effects of dispersal date on winter flock establishment and social dominance in marsh tits *Parus palustris*. Journal of Animal Ecology, 57(3), 917–928. doi: 10.2307/5101

Norris, D. R., & Marra, P. P. (2007). Seasonal interactions, habitat quality, and population dynamics in migratory birds. The Condor, 109(3), 535–547. doi: 10.1650/8350.1

Norris, D. R., Marra, P. P., Kyser, T. K., Sherry, T. W., & Ratcliffe, L. M. (2004). Tropical winter habitat limits reproductive success on the temperate breeding grounds in a migratory bird. Proceedings of the Royal Society B: Biological Sciences, 271(1534), 59–64. doi: 10.1098/rspb.2003.2569

Nussey, D. H., Postma, E., Gienapp, P., & Visser, M. E. (2005). Selection on heritable phenotypic plasticity in a wild bird population. Science, 310(5746), 304–306. doi: 10.1126/science.1117004

Odum, E. P. (1941a). Winter homing behavior of the chickadee. Bird-banding, 12(3), 113–119.

Odum, E. P. (1941b). Annual cycle of the black-capped chickadee: 1. The Auk, 58(3), 314–333. doi: 10.2307/4078950

Ortego, J., García-Navas, V., Ferrer, E. S., & Sanz, J. J. (2011). Genetic structure reflects natal dispersal movements at different spatial scales in the blue tit, *Cyanistes caeruleus*. Animal Behaviour, 82(1), 131–137. doi: 10.1016/j.anbehav.2011.04.007

Pakanen, V. -M., Koivula, K., Orell, M., Rytkönen, S., & Lahti, K. (2016). Sex-specific mortality costs of dispersal during the post-settlement stage promote male philopatry in a resident passerine. Behavioral Ecology and Sociobiology, 70(10), 1727–1733. doi: 10.1007/s00265-016-2178-z

Perrins, C. M. (1970). The timing of birds’ breeding seasons. Ibis, 112(2), 242–255. doi: 10.1111/j.1474-919X.1970.tb00096.x

Perrins, C. M. (1979). British Tits. London, U.K.: Collins.

Perrins, C. M., & McCleery, R. H. (1989). Laying dates and clutch size in the great tit. The Wilson Bulletin, 101(2), 236–253.

Petersen, M. R. (1992). Reproductive ecology of emperor geese: annual and individual variation in nesting. The Condor, 94(2), 383–397. doi: 10.2307/1369211

Piersma, T. (1987). Hop, skip or jump? Constraints on migration of Arctic waders by feeding, fattening, and flight speed. Limosa, 60, 185–194.

Psorakis, I., Roberts, S. J., Rezek, I., & Sheldon, B. C. (2012). Inferring social network structure in ecological systems from spatio-temporal data streams. Journal of the Royal Society Interface, 9(76), 3055–3066. doi: 10.1098/rsif.2012.0223

R Core Team. (2017). R: A language and environment for statistical computing. R Foundation for Statistical Computing, Vienna, Austria. URL https://www.R-project.org/.

Rainio, K., Tøttrup, A. P., Lehikoinen, E., & Coppack, T. (2007). Effects of climate change on the degree of protandry in migratory songbirds. Climate Research, 35, 107–114. doi: 10.3354/cr00717

Saino, N., Ambrosini, R., Caprioli, M., Romano, A., Romano, M., Rubolini, D., … Liechti, F. (2017). Sex-dependent carry-over effects on timing of reproduction and fecundity of a migratory bird. Journal of Animal Ecology, 86(2), 239–249. doi: 10.1111/1365-2656.12625

Sandell, M., & Smith, H. G. (1991). Dominance, prior occupancy, and winter residency in the great tit (*Parus major*). Behavioral Ecology and Sociobiology, 29(2), 147–152. doi: 10.1007/BF00166490

Schlicht, E., & Kempenaers, B. (2015). Immediate effects of capture on nest visits of breeding blue tits, *Cyanistes caeruleus*, are substantial. Animal Behaviour, 105, 63–78. doi: 10.1016/j.anbehav.2015.04.010

Schlicht, L., Girg, A., Loës, P., Valcu, M., & Kempenaers, B. (2012). Male extrapair nestlings fledge first. Animal Behaviour, 83(6), 1335–1343. doi: 10.1016/j.anbehav.2012.02.021

Schlicht, L., Valcu, M., & Kempenaers, B. (2015). Spatial patterns of extra-pair paternity: beyond paternity gains and losses. Journal of Animal Ecology, 84(2), 518–531. doi: 10.1111/1365-2656.12293

Silk, M. J., Croft, D. P., Tregenza, T., & Bearhop, S. (2014). The importance of fission-fusion social group dynamics in birds. Ibis, 156(4), 701–715. doi: 10.1111/ibi.12191

Smallegange, I. M., Fiedler, W., Köppen, U., Geiter, O., & Bairlein, F. (2010). Tits on the move: exploring the impact of environmental change on blue tit and great tit migration distance. Journal of Animal Ecology, 79(2), 350–357. doi: 10.1111/j.1365-2656.2009.01643.x

Smith, S. M. (1995). Age-specific survival in breeding black-capped chickadees (*Parus atricapillus*). The Auk, 112(4), 840–846. doi: 10.2307/4089016

Smith, H. G., & Nilsson, J. (1987). Intraspecific variation in migratory pattern of a partial migrant, the blue tit (*Parus caeruleus*): an evaluation of different hypotheses. The Auk, 104(1), 109–115. doi: 10.2307/4087239

Smith, R. J., & Moore, F. R. (2003). Arrival fat and reproductive performance in a long-distance passerine migrant. Oecologia, 134(3), 325–331. doi: 10.1007/s00442-002-1152-9

Smith, R. J., & Moore, F. R. (2005). Arrival timing and seasonal reproductive performance in a long-distance migratory landbird. Behavioral Ecology and Sociobiology, 57(3), 231–239. doi: 10.1007/s00265-004-0855-9

Stanley, C. Q., MacPherson, M., Fraser, K. C., McKinnon, E. A., & Stutchbury, B. J. M. (2012). Repeat tracking of individual songbirds reveals consistent migration timing but flexibility in route. PloS One, 7(7), e40688. doi: 10.1371/journal.pone.0040688

Stenning, M. (2018). The Blue Tit. Bloomsbury Publishing.

Stoffel, M. A., Nakagawa, S., & Schielzeth, H. (2017). rptR: repeatability estimation and variance decomposition by generalized linear mixed-effects models. Methods in Ecology and Evolution, 8(11), 1639–1644. doi: 10.1111/2041-210X.12797

Tarka, M., Hansson, B., & Hasselquist, D. (2015). Selection and evolutionary potential of spring arrival phenology in males and females of a migratory songbird. Journal of Evolutionary Biology, 28(5), 1024–1038. doi: 10.1111/jeb.12638

Thorley, J. B., & Lord, A. M. (2015). Laying date is a plastic and repeatable trait in a population of blue tits *Cyanistes caeruleus*. Ardea, 103(1), 69–78. doi: 10.5253/arde.v103i1.a7

Tryjanowski, P., Sparks, T. H., Ptaszyk, J., & Kosicki, J. (2004). Do white storks *Ciconia ciconia* always profit from an early return to their breeding grounds? Bird Study, 51(3): 222–227. doi: 10.1080/00063650409461357

Valcu, M., & Kempenaers, B. (2008). Causes and consequences of breeding dispersal and divorce in a blue tit, *Cyanistes caeruleus*, population. Animal Behaviour, 75(6), 1949–1963. doi: 10.1016/j.anbehav.2007.12.005

van Wijk, R. E., Bauer, S., & Schaub, M. (2016). Repeatability of individual migration routes, wintering sites, and timing in a long-distance migrant bird. Ecology and Evolution, 6(24), 8679–8685. doi: 10.1002/ece3.2578

Vardanis, Y., Klaassen, R. H. G., Strandberg, R., & Alerstam, T. (2011). Individuality in bird migration: Routes and timing. Biology Letters, 7(4), 502–505. doi: 10.1098/rsbl.2010.1180

Vergara, P., Aguirre, J. I., & Fernández-Cruz, M. (2007). Arrival date, age and breeding success in white stork *Ciconia ciconia*. Journal of Avian Biology, 38(5), 573–579. doi: 10.1111/j.2007.0908-8857.03983.x

Verhulst, S., & Nilsson, J. A. (2008). The timing of birds’ breeding seasons: a review of experiments that manipulated timing of breeding. Philosophical Transactions of the Royal Society B: Biological Sciences, 363(1490), 399–410. doi: 10.1098/rstb.2007.2146

Verhulst, S., Perrins, C. M., & Riddington, R. (1997). Natal dispersal of great tits in a patchy environment. Ecology, 78(3), 864–872. doi: 10.1890/0012-9658(1997)078[0864:NDOGTI]2.0.CO;2

Verhulst, S., & Tinbergen, J. M. (1991). Experimental evidence for a causal relationship between timing and success of reproduction in the great tit *Parus m. major*. Journal of Animal Ecology, 60(1), 269–282. doi: 10.2307/5459

Village, A. (1985). Spring arrival times and assortative mating of kestrels in south Scotland. Journal of Animal Ecology, 54(3), 857–868. doi: 10.2307/4383

Williams, T. D. (2012). Physiological adaptations for breeding in birds. Princeton (NJ): Princeton University Press.

Wilson, A., & Nussey, D. H. (2010). What is individual quality? An evolutionary perspective. Trends in Ecology and Evolution, 25(4), 207–214. doi: 10.1016/j.tree.2009.10.002

